# Two pathways of p27^Kip1^ degradation are required for murine lymphoma driven by Myc and EBV latent membrane protein 2A

**DOI:** 10.1101/564559

**Authors:** Richard P. Sora, Masato Ikeda, Richard Longnecker

**Author notes:** **Corresponding Author:** Richard Longnecker. **Competing Interests:** Authors declare no competing financial interests.

## Abstract

Epstein-Barr virus (EBV) latent membrane protein 2A (LMP2A), expressed in EBV latency, contributes to Burkitt Lymphoma (BL) development in a murine model by acting as a constitutively active B cell receptor (BCR) mimic. Mice expressing both LMP2A and *MYC* transgenes (LMP2A/λ-*MYC*) develop tumors significantly faster than mice only expressing *MYC* (λ-*MYC*). Previously, we demonstrated the cell cycle inhibitor p27^Kip1^ is present at significantly lower levels in LMP2A/λ-*MYC* mice due to increased post-translational degradation. P27^Kip1^ degradation can occur in the cytoplasm following phosphorylation on serine 10 (S10), or in the nucleus via the SCF^Skp2^ complex, which depends on Cks1. We previously demonstrated a S10A knock-in of p27^Kip1^ (p27^S10A/S10A^), which prevented S10 phosphorylation, failed to significantly delay tumor onset in LMP2A/λ-*MYC* mice. We also previously demonstrated that a *Cks1* knockout partially delayed tumor onset in LMP2A/λ-*MYC* mice, but onset was still significantly faster than in λ-*MYC* mice. Here, we have combined both genetic manipulations in what we call p27^Super^ mice. LMP2A/λ-*MYC*/p27^Super^ mice and λ-*MYC*/p27^Super^ mice both displayed dramatic delays in tumor onset. Strikingly, tumor development in LMP2A/λ-*MYC*/p27^Super^ mice was later than in λ-*MYC* mice and not significantly different from λ-*MYC*/p27^Super^ mice. The p27^Super^ genotype also normalized G_1_-S phase cell cycle progression, spleen size, and splenic architecture in LMP2A/λ-*MYC* mice. Our results reveal both major pathways of p27^Kip1^ degradation are required for the accelerated BL development driven by LMP2A in our BL model and that blocking both degradation pathways is sufficient to delay Myc-driven tumor development with or without LMP2A.

**Importance:** Burkitt lymphoma (BL) is a cancer that primarily affects children. The side effects of chemotherapy highlight the need for better BL treatments. Many BL tumors contain Epstein-Barr virus (EBV) and our goal is to determine what makes EBV-positive BL different from EBV-negative BL. This may lead to more specific treatments for both types. All cases of BL require overexpression of *MYC*. Mice engineered to express an EBV LMP2A along with *MYC* (LMP2A/λ-*MYC* mice) develop tumors much more quickly than mice only expressing MYC (λ-*MYC* mice). Blocking degradation of the cell cycle inhibitor protein p27^Kip1^ in LMP2A/λ-*MYC* mice causes tumors to develop later than in λ-*MYC* mice, showing that p27^Kip1^ degradation may play a larger role in EBV-positive BL than EBV-negative. Furthermore, our studies suggest the cell cycle may be an attractive target as a treatment option for LMP2A positive cancers in humans.

## Introduction

Epstein-Barr virus (EBV) is a γ-herpesvirus that establishes latent infection in over 95% of the world’s population by adulthood(1). EBV latency occurs primarily in B cells and has been associated with several B cell cancers, including Burkitt Lymphoma (BL)(2). There are three recognized BL subtypes: endemic (eBL), found mainly in equatorial Africa, sporadic (sBL), found throughout the rest of the world, and AIDS-associated (AIDS-BL), found in HIV-positive patients. Nearly all eBL tumors, and smaller proportions of sBL and AIDS-BL tumors, are associated with EBV(1). *In vitro*, EBV can transform resting B cells into growing lymphoblastoid cell lines by expressing the “growth program”, which drives cell proliferation and survival(3–5). *In vivo*, however, EBV exploits B cell biology to achieve latency.

In order for naïve B cells to become memory B cells, they undergo clonal expansion following antigen exposure and form germinal centers (GC) in the spleen and lymph nodes(5, 6). While in the GC, antigen binding to the B cell receptor (BCR) leads to phosphorylation of the tyrosine kinase Syk, which drives pro-survival signaling through proteins such as PI3K, Akt, and ERK(7). The clones that survive begin to differentiate and exit the GC, becoming memory B cells(7). Taking advantage of this process, EBV infects naïve B cells, driving clonal expansion of EBV-infected B cells(5). Once in the GC, EBV transitions to a latency program in which a small number of genes including Latent Membrane Proteins 1 (LMP1) and 2A (LMP2A), as well as EBNA1 are expressed(5). In this environment, LMP2A acts as a constitutively active mimic of the BCR, driving survival and differentiation(8, 9). As the infected B cells differentiate into memory B cells, EBV switches to another latency program, expressing EBNA1 and other non-protein-coding genes(5, 10). B cells isolated from BL tumors display a GC phenotype and, while EBV-positive BL tumors were first observed expressing only EBNA1, LMP2A transcripts were subsequently detected in eBL cell lines and snap-frozen eBL tumors(11–17). It is therefore likely that EBV-infected B cells transform into BL tumor cells when EBV drives clonal proliferation early after EBV infection.

A key feature of all BL tumors is a translocation of the proto-oncogene c-*MYC* (*MYC*) and an immunoglobulin (Ig) gene locus, leading to Myc overexpression under an Ig promoter(18). A C57BL/6 murine model of BL has been developed in which a *MYC* transgene is expressed under the Igλ locus (λ-*MYC*)(19). Myc is a transcription factor that drives many types of cancer. It promotes the expression of genes that promote cell cycle progression from G_1_ to S phase and is required for normal B cell activation and proliferation(20). In addition, Myc induces expression of several tumor suppressor genes that promote cell cycle arrest and apoptosis, including *TP53* and *ARF*(21). For this reason, inactivation of tumor suppressor pathways is required for the development of Myc-driven tumors. When λ-*MYC* mice are crossed with LMP2A transgenic mice, the resulting LMP2A/λ-*MYC* mice develop tumors significantly faster than λ-*MYC* mice (22, 23). We previously found that spleens of LMP2A/λ-*MYC* mice displayed a greater percentage of S-phase B cells than λ-*MYC* mice and the cell cycle inhibitor p27^Kip1^ was rapidly degraded and expressed at lower levels in LMP2A/λ-*MYC* splenic B cells(24).

Degradation of p27^Kip1^ occurs through two major pathways. Phosphorylation on serine 10 (S10), results in the export from the nucleus and subsequent degradation of p27^Kip1^ in the cytoplasm(25). Alternatively, phosphorylation can occur on threonine 187 (T187) leading to degradation of p27^Kip1^ by the SCF^Skp2^ complex in the nucleus(26). Previously, we attempted to delay lymphomagenesis in LMP2A/λ-*MYC* mice by using a p27^Kip1^ S10A mutant homozygous knock-in (p27^S10A/S10A^) mouse, which proved unsuccessful, although we did observe modest effects on G_1_-S cell cycle progression in λ-*MYC* mice(24). Subsequent research determined that preventing p27^Kip1^ degradation by SCF^Skp2^ via the homozygous knockout of *Cks1* (*Cks1*^−/−^), a member of the SCF^Skp2^ complex that is essential for the recognition of phosphorylated T187, resulted in a significant delay in lymphomagenesis for LMP2A/λ-*MYC* mice(27). These mice, however, still developed tumors much faster than the λ-*MYC* mice(27).

To investigate the two pathways of p27^Kip1^ degradation in LMP2A-mediated lymphomagenesis, we crossed *Cks1* knockout mice (8) with p27^S10A/S10A^ mice (28) and identified offspring that were homozygous for the S10A knock-in and *Cks1* knockout, which we termed p27^Super^ mice. Both LMP2A/λ-*MYC*/p27^Super^ and λ-*MYC*/p27^Super^ mice displayed dramatically delayed tumor onset. Strikingly, tumor onset in the LMP2A/λ-*MYC*/p27^Super^ mice was later than in the λ-*MYC* mice and not significantly different from λ-*MYC*/p27^Super^ mice. These data show that preventing p27^Kip1^ degradation by using the p27^Super^ genotype is the most effective genetic manipulation yet for preventing tumor growth in our model of BL.

## Results

### The p27^Super^ genotype delays tumor onset in both LMP2A/λ-*MYC* and λ-*MYC* mice

To observe the effect of degradation-resistant p27^Kip1^ on Myc-induced tumorigenesis, tumor-free survival in LMP2A/λ-*MYC*, LMP2A/λ-*MYC*/p27^Super^, λ-*MYC*, and λ-*MYC*/p27^Super^ mice was examined (Fig. 1). All tumors appeared in lymph nodes in the cervical, abdominal, or thoracic area. Tumor onset was delayed in both LMP2A/λ-*MYC*/p27^Super^ and λ-*MYC*/p27^Super^ mice (Fig. 1). Median tumor-free survival time was 378 days in LMP2A/λ-*MYC*/p27^Super^ mice compared to 63 in LMP2A/λ-*MYC* mice, a delay of 315 days. In our previous study, median tumor-free survival in LMP2A/λ-*MYC*/*Cks1*^−/−^ mice was delayed only 61.5 days compared to LMP2A/λ-*MYC*(27). The S10A knock-in alone was previously shown to have no effect on tumor onset (36 days for both the LMP2A/λ-*MYC*/p27^S10A/S10A^ and LMP2A/λ-*MYC* mice)(24). Strikingly, tumor onset in the LMP2A/λ-*MYC*/p27^Super^ mice was 172 days later than in the λ-*MYC* mice, and there was no significant difference in tumor onset between the LMP2A/λ-*MYC*/p27^Super^ and the λ-*MYC*/p27^Super^ mice, which had a median tumor-free survival time of 400 days. In our previous study, tumor onset in LMP2A/λ-*MYC*/*Cks1*^−/−^ mice was 25 days earlier than λ-*MYC* mice(27). Although the p27^S10A/S10A^ knock-in by itself was previously shown to have no effect on tumor-free survival(24), our data show that it improved tumor-free survival in the presence of the *Cks1* knockout. From this data, we conclude that blocking both major pathways of p27^Kip1^ degradation completely blocked accelerated LMP2A-driven lymphomagenesis in our BL model.

**Figure 1:**
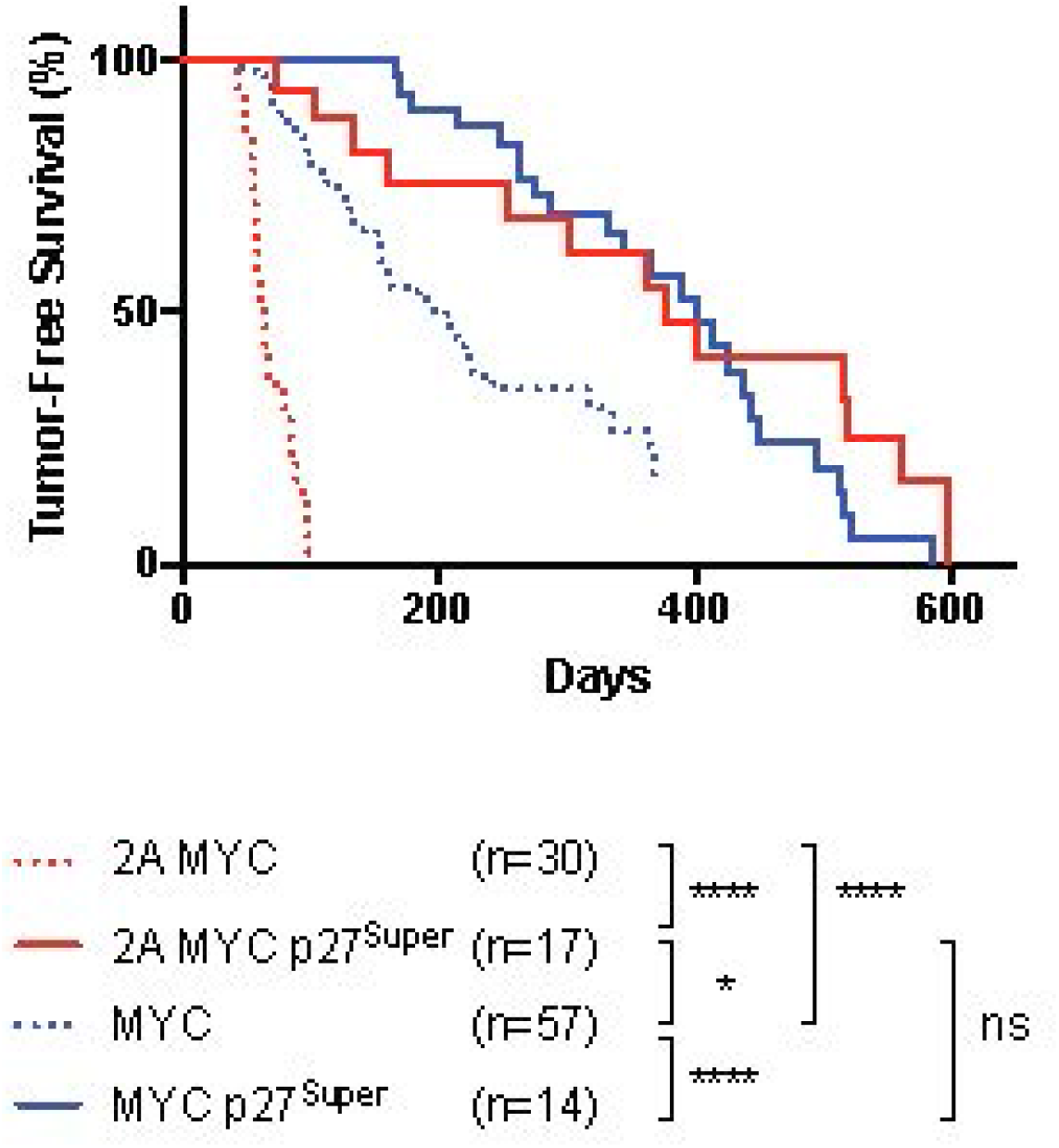
The p27^Super^ genotype delays MYC-driven tumor development and blocks the accelerated tumor onset driven by LMP2A. Tumor-free survival curve showing the number of days for discernible tumors to develop in the cervical, abdominal, or thoracic area for each of the four tumor-associated genotypes. Tumor onset was delayed 315 days in LMP2A/λ-*MYC*/p27^Super^ compared to LMP2A/λ-*MYC* mice. In previous studies, tumor onset was delayed 61.5 days in LMP2A/λ-*MYC*/*Cks1*^−/−^ and not delayed in the LMP2A/λ-*MYC*/p27^S10A/S10A^ mice (24, 27). Tumor onset in LMP2A/λ-*MYC*/p27^Super^ mice is 172 days later than λ-*MYC* mice. Previously, tumor onset in LMP2A/λ-*MYC*/*Cks1*^−/−^ mice was 25 days earlier than λ-*MYC* mice(27). Sample size (n) for each genotype is indicated below the curve, as well as p values determined by log-rank (Mantel-Cox) test. * p<0.05, **** p<0.0001, ns: not significant.

### The p27^Super^ genotype does not alter splenic cellular architecture or B cell numbers in the spleen compared to WT

We next investigated whether the p27^Super^ genotype affects normal B cell development in 4 week old mice. The p27^Super^ mice developed a similar number of mature B cells in the periphery (Fig 2A). B cell numbers were lower in the bone marrow of p27^Super^ mice, whereas the splenic B cell number in p27^Super^ mice was similar to WT mice (Fig. 2A). We next analyzed splenic architecture in p27^Super^ mice compared to WT mice by staining spleens for B220 and p27^Kip1^. B cell development is a highly regulated process, which results in the formation of follicles of developing B cells that can be readily observed in the spleen by immunohistochemistry (IHC). B cell follicle formation and p27^Kip1^ staining were similar in the p27^Super^ spleens compared to WT spleens (Fig. 2B), demonstrating that the p27^Super^ genotype does not significantly alter splenic B cell number or follicle formation.

**Figure 2:**
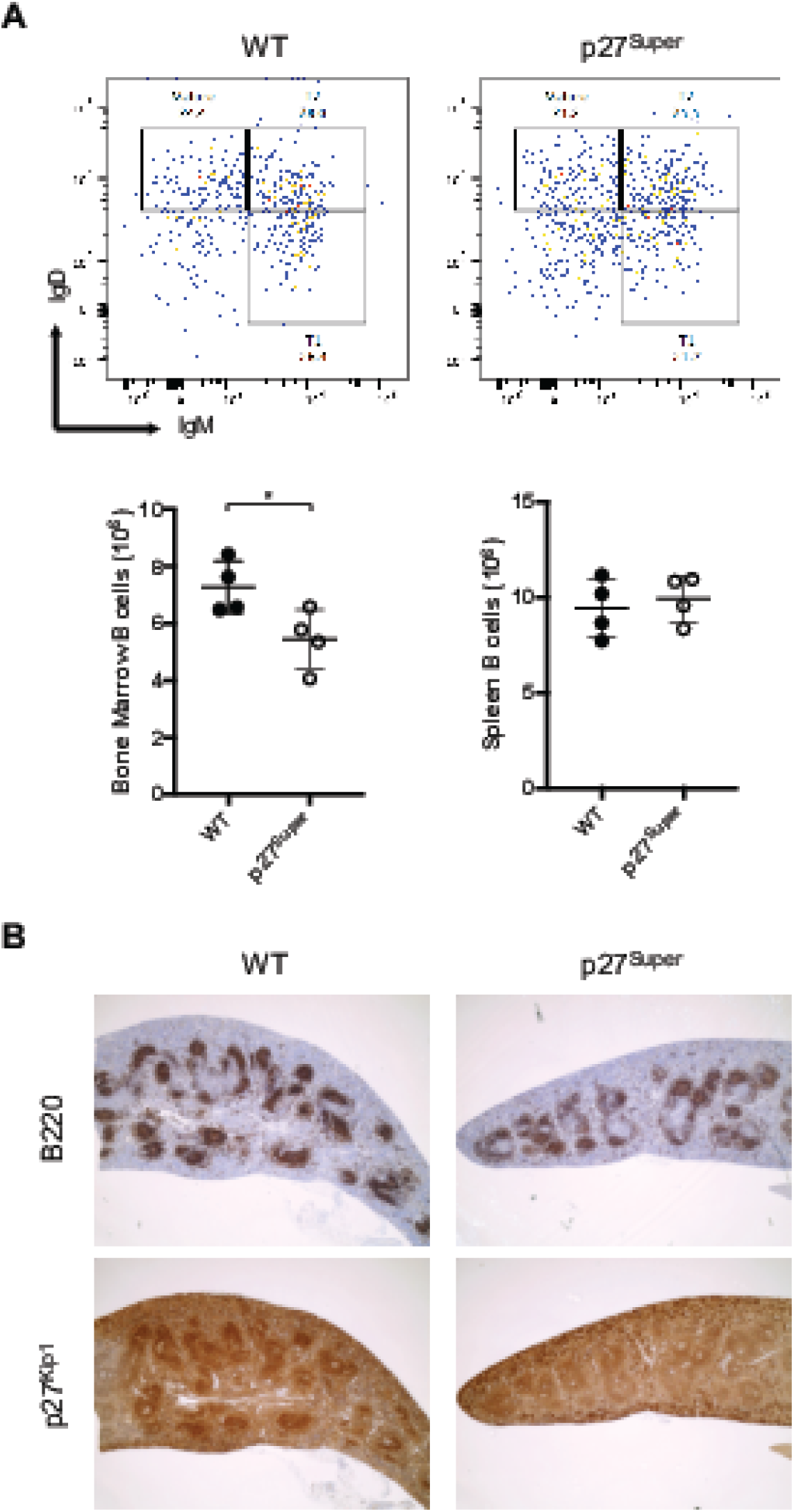
Number and follicular formation of pretumor splenic B cells are not altered in p27^Super^ mice. (A) Total B cell number in bone marrow and spleens of WT and p27^Super^ mice determined by flow cytometry. Spleens were dissected from 4-5 week-old mice of WT and p27^Super^ genotypes. B220^+^ B cells were plotted with IgM and IgD. T1: IgM^high^ IgD^low^ transitional B cells. T2: IgM^high^ IgD^high^ transitional B cells. Mature: Mature B cells. (B) Immunohistochemistry (IHC) of 6 to 8-week old mouse spleens stained with B220 and p27^Kip1^ and imaged with 4X magnification. Data represent the mean +/− SD. P values were determined by two-tailed t-test. * p<0.05.

### The p27^Super^ genotype normalizes B cell development in LMP2A/λ-*MYC* and λ-*MYC* spleen

Our previous studies showed pretumor LMP2A/λ-*MYC* and λ-*MYC* mice have a significantly higher percentage of splenic B cells in S phase than WT mice of the same age (24, 27). By analyzing the cell cycle, we found the percentage of S phase cells in LMP2A/λ-*MYC*/p27^Super^ spleens was 27.7% lower than in LMP2A/λ-*MYC* mice and similar to what is observed in WT mice (Fig. 3A). In our previous studies, the percentage of pretumor splenic B cells in S-phase was 0.59% lower in LMP2A/λ-*MYC*/p27^S10A/S10A^ mice and 19.8% lower in LMP2A/λ-*MYC*/*Cks1*^−/−^ mice than in LMP2A/λ-*MYC*(24, 27). The percentage of splenic S phase B cells was 10.5% lower in λ-*MYC*/p27^Super^ spleens than λ-*MYC* (Fig. 3A), while in previous studies the percentage was 9.13% lower in λ-MYC/p27^S10A/S10A^ and 6.5% lower in λ-*MYC*/*Cks1*^−/−^ mice(24, 27) compared to λ-*MYC* mice. Finally, the percentages of S phase cells in LMP2A/λ-*MYC*/p27^Super^ and λ-*MYC*/p27^Super^ mice were not significantly different (18% and 20% respectively, Fig 3A), compatible with the similar tumor onset observed in these two genotypes. These data show that blocking both p27^Kip1^ degradation pathways together has a synergistic effect on halting G_1_-S phase cell cycle transition, and that this synergy is specific to LMP2A/λ-*MYC* mice.

**Figure 3:**
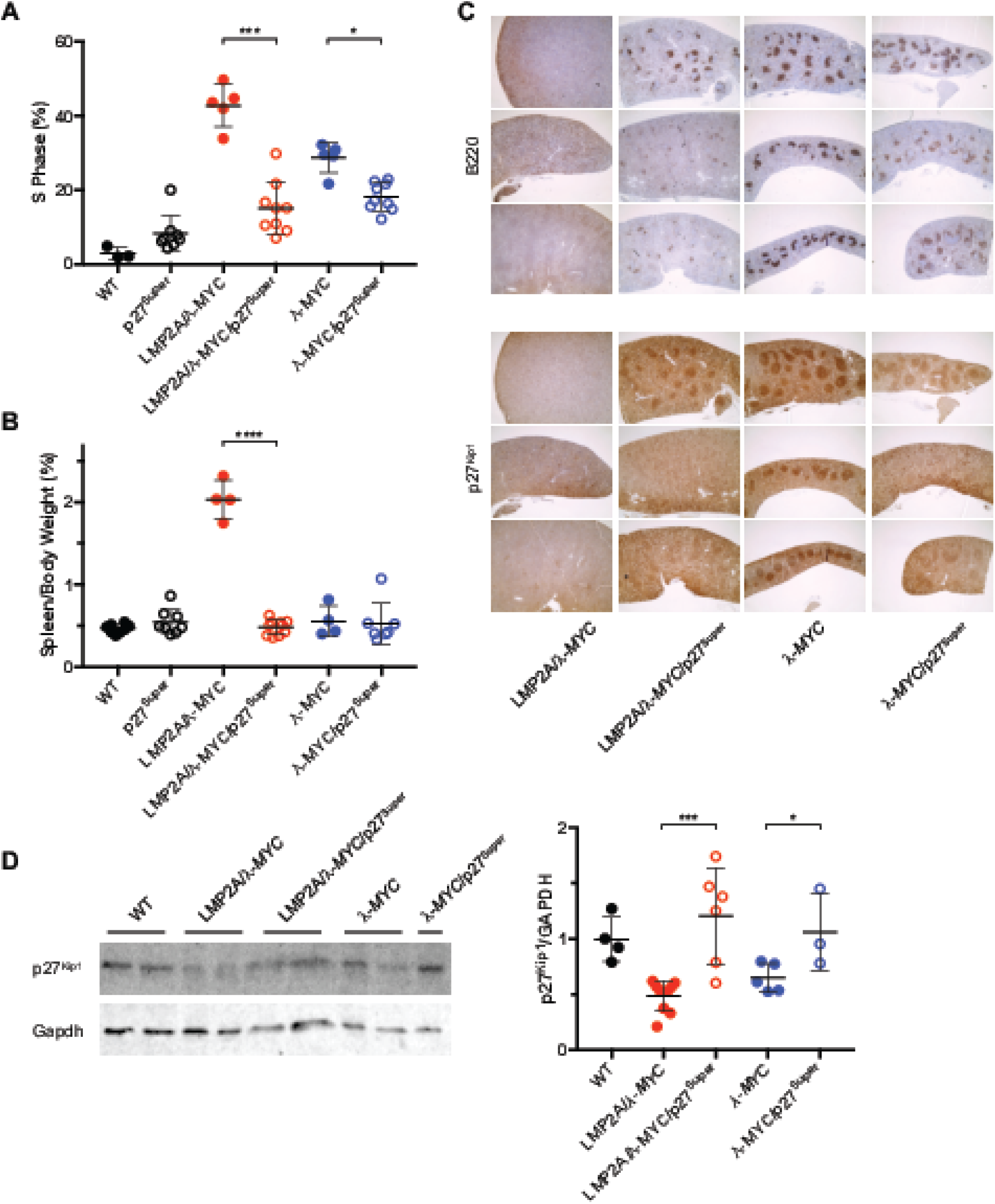
λ-*MYC*/p27^Super^ and LMP2A/λ-*MYC*/p27^Super^ mice display normal spleen size and B cell development, as well as elevated p27^Kip1^ levels in pretumor splenic B cells. (A) Cell cycle analysis of WT, p27^Super^, LMP2A/λ-*MYC*, LMP2A/λ-*MYC*/p27^Super^, λ-*MYC*, λ-*MYC*/p27^Super^ pretumor splenic B cells. Graph represents the percentage of splenic B cells that are in S phase. The percentage of splenic B cells in S phase is 27.7% lower in LMP2A/λ-*MYC*/p27^Super^ mice than LMP2A/λ-*MYC* mice, a greater difference than what was previously observed in LMP2A/λ-*MYC*/*Cks1*^−/−^ (19.8%) and LMP2A/λ-*MYC*/p27^S10A/S10A^ (0.59%)(24, 27) (B) The ratios of spleen weight to total body weight were calculated for 4 to 5-week old mice of the indicated genotypes. (C) IHC was performed on spleens of 6 to 8-week old pretumor mice as described in Figure 1. Spleens were sectioned and stained for B220 and p27^Kip1^. Three mouse samples are shown for each genotype. (D) Immunoblots were performed on protein isolated from splenic B cells of pretumor mice. Membranes were probed for p27^Kip1^ and GAPDH. Densitometry was performed to calculate relative p27^Kip1^ levels. Representative blot (Left) and relative p27^Kip1^ levels (Right) are shown. Data represent the mean +/− SD. P values were determined by twotailed t-test. * p<0.05, *** p<0.001, **** p<0.0001.

The p27^Super^ genotype also prevented the splenomegaly observed in LMP2A/λ-*MYC* mice. As previously shown(24, 27), LMP2A/λ-*MYC* mice at 4-5 weeks old displayed significantly higher spleen weight relative to body weight than WT mice (Fig. 3B). The LMP2A/λ-*MYC*/p27^Super^ mice, however, had spleen-to-body weight ratios that were not significantly different from WT mice (Fig. 3B).

After exiting the bone marrow, normal B cells form follicles in the spleen and other secondary lymphoid organs. This B cell organization is completely lost in the spleens of LMP2A/λ-*MYC* mice (Fig. 3C). This suggests that the LMP2A/λ-*MYC* genotype leads to a defect of normal B cell development or follicular homing. The p27^Super^ genotype, however, restored B cell follicle formation in the LMP2A/λ-*MYC*/p27^Super^ mice (Fig. 3C). While the B cell follicles in all four λ-*MYC* genotype-containing spleens appeared to be less defined than in WT, B cell organization in LMP2A/λ-*MYC*/p27^Super^, *λ-MYC*, and λ-*MYC*/p27^Super^ spleens closely resembled WT rather than the disorganized LMP2A/λ-*MYC* spleens (Fig. 3C).

### P27^Kip1^ protein degradation is blocked in p27^Super^ mice

We next analyzed p27^Kip1^ levels in splenic B cells. Immunohistostaining of pretumor spleens showed an observable increase in the intensity of the p27^Kip1^ signal in both the LMP2A/λ-*MYC*/p27^Super^ and λ-*MYC*/p27^Super^ mice compared to LMP2A/λ-*MYC* and λ-*MYC* mice, respectively (Fig. 3C). To confirm the IHC data, we performed western blot analysis to quantify p27^Kip1^ levels in pretumor splenic B cells (Fig. 3D). Spleens were dissected from mice prior to tumor onset and B cells were isolated and purified. As expected, p27^Kip1^ levels were significantly elevated in LMP2A/λ-*MYC*/p27^Super^ and λ-*MYC*/p27^Super^ mice compared to LMP2A/λ-*MYC* and λ-*MYC* mice, respectively. In our previous studies, the *Cks1* knockout alone increased p27^Kip1^ levels in LMP2A/λ-*MYC* nearly to wild type levels, but resulted in only a partial delay in tumor development(27). In our current study, the LMP2A/λ-*MYC*/p27^Super^ and λ-*MYC*/p27^Super^ mice have higher p27^Kip1^ levels than WT mice, although the difference is not statistically significant. This greater increase in p27^Kip1^ levels may account for the greater delay in tumor onset caused by the p27^Super^ genotype than by the *Cks1* knockout alone.

### The p27^Super^ genotype significantly increases p27^Kip1^ levels in both LMP2A/λ-*MYC* and λ-*MYC* tumors

In order to determine whether p27^Kip1^ levels remain elevated in LMP2A/λ-*MYC*/p27^Super^ and λ-*MYC*/p27^Super^ tumors, mice were sacrificed once tumors became apparent. Lymph node tumors were dissected from either the cervical or abdominal area. IHC was performed to assess p27^Kip1^ levels (Fig. 4A) as was done with pretumor spleens. Western blots were also performed to quantify the levels of p27^Kip1^ in B cells isolated from the tumors (Fig. 4B). Both the IHC and Western blot analysis showed that p27^Kip1^ levels are significantly elevated in LMP2A/λ-*MYC*/p27^Super^ and λ-*MYC*/p27^Super^ tumors compared to LMP2A/λ-*MYC* and λ-MYC, respectively. These data suggest that while tumors still form in LMP2A/λ-*MYC*/p27^Super^ and λ-*MYC*/p27^Super^ mice, it is likely due to factors other than p27^Kip1^ degradation.

**Figure 4:**
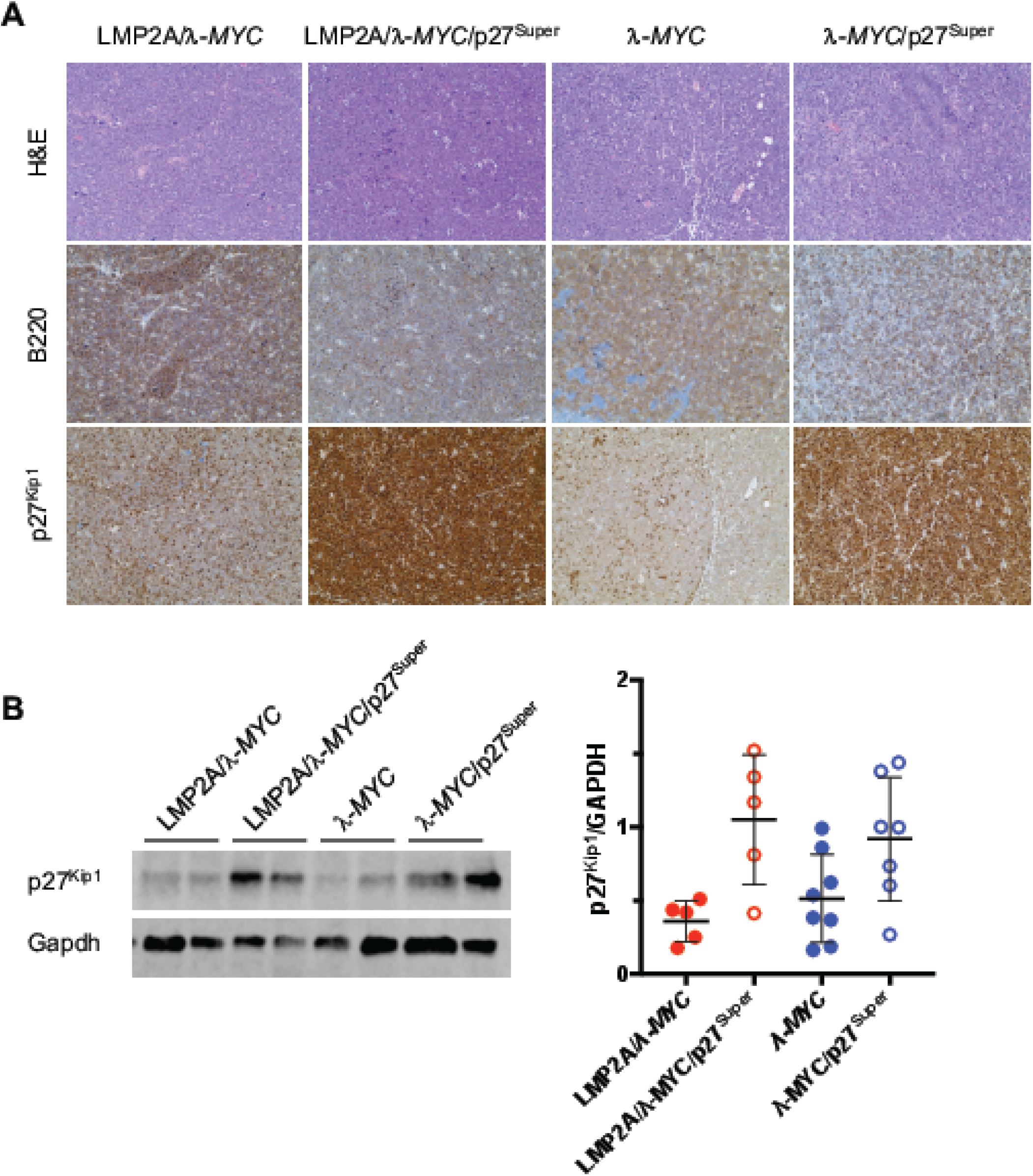
Tumors from λ-*MYC*/p27^Super^ and LMP2A/λ-*MYC*/p27^Super^ mice maintain elevated p27^Kip1^ levels. (A) IHC was performed on lymph node tumors dissected from mice of the indicated genotypes stained with H&E, B220 and p27^Kip1^ and imaged with ×40 magnification microscope. (B) Immunoblots were performed on protein isolated from mouse tumors of the indicated genotypes. Representative blot (Left) and relative p27^Kip1^ levels (Right) are shown. Blots and statistics were performed as described in Figure 3. Data represent the mean +/− SD. * p<0.05.

## Discussion

The mechanisms driving BL tumors can differ drastically depending on the presence or absence of EBV. EBV-positive BL genomes carry fewer, and distinct, driver mutations compared to EBV-negative(29). Previous studies, including by our group, have shown that LMP2A accelerates tumor development in combination with dysregulated Myc, obviating the need for mutations in the ARF-Mdm2-p53 pathway(22, 23). We have also demonstrated that LMP2A cooperates with Myc to increase G_1_-S phase cell cycle transition prior to tumorigenesis(24, 27). Determining the effect of EBV latent proteins like LMP2A on the cell cycle can uncover important distinctions between EBV-positive and EBV-negative BL, which may lead to more specific therapies for BL patients based on disease subtype.

Gene expression profiling has previously shown elevated expression of genes involved in the cell cycle regulation in eBL compared to sBL(30). Additionally, RNA-seq found mutations in *CCND3*, which encodes cyclin D3 a critical regulator of G_1_-S transition, were frequent in sBL but not eBL(31). Furthermore, expression of LMP2A in eBL correlates with increased expression of genes promoting G_1_-S phase cell cycle transition(32). In our previous studies, post-translational degradation of p27^Kip1^ correlated with earlier tumor development in LMP2A/λ-*MYC* mice, which was partially attenuated by blocking Cks1-dependent p27^Kip1^ degradation(27). As noted in the Introduction, our previous attempts to delay tumor onset in LMP2A/λ-*MYC* and λ-*MYC* mice by blocking S10 phosphorylation alone were unsuccessful. In the current study, we delayed tumor onset by blocking both Cks1-dependent and S10 phosphorylation-dependent p27^Kip1^ degradation with the p27^Super^ genotype. We found that p27^Super^ mice had a dramatic delay in tumor onset when compared to the *Cks1* knockout alone in both LMP2A/λ-*MYC*/p27^Super^ mice and λ-*MYC*/p27^Super^ mice.

Of twelve proteins known to regulate G_1_-S phase transition (p27^Kip1^, p21^Cip1^, p15^Ink4b^, p16^Ink4a^, Cyclin D1, D2, D3, E and A, CDK2, Rb and E2f1), p27^Kip1^ was the only one expressed at significantly lower levels in LMP2A/λ-*MYC* than in λ-*MYC* pretumor B cells(24). For this reason, we predicted that blocking p27^Kip1^ degradation in LMP2A/λ-*MYC* mice would prevent tumors from developing more quickly than in λ-*MYC* mice. Our results show that tumor onset in LMP2A/λ-*MYC*/p27^Super^ mice was significantly later than in λ-*MYC* mice, and not significantly different from λ-*MYC*/p27^Super^ mice. With the p27^Super^ genotype, mice expressing LMP2A and Myc do not develop tumors more quickly than mice only expressing Myc, demonstrating that both pathways of p27^Kip1^ degradation are required for the accelerated tumorigenesis driven by LMP2A in the presence of dysregulated Myc.

The effect of the p27^Super^ genotype on the pretumor spleen and splenic B cells mirrored its effect on tumor onset. The percentage of splenic B cells in S phase was significantly decreased in LMP2A/λ-*MYC*/p27^Super^ and λ-*MYC*/p27^Super^ mice (Fig. 3A) and the splenomegaly observed in LMP2A/λ-*MYC* mice was normalized in LMP2A/λ-*MYC*/p27^Super^ mice (Fig. 3B). Additionally, B cell follicle formation is partially restored in LMP2A/λ-*MYC*/p27^Super^ pretumor spleens (Fig. 3C). The increased p27^Kip1^ expression observed in LMP2A/λ-*MYC*/p27^Super^ and λ-*MYC*/p27^Super^ splenic B cells (Fig. 3D) is maintained in tumors from both genotypes (Fig 4B), suggesting these tumors develop independently of p27^Kip1^ degradation. Interestingly, in the absence of λ-*MYC*, p27^Super^ does not affect splenic B cell number (Fig. 1A) and does not decrease the percentage of S-phase B cells (Fig. 3A) or spleen-to-body ratio (Fig 3B). This suggests the effects of p27^Super^ on the cell cycle and B cell development are specific to LMP2A/λ-*MYC* and λ-*MYC* mice.

While there is no significant difference in tumor onset between the LMP2A/λ-*MYC*/p27^Super^ and λ-*MYC*/p27^Super^ mice, the pathways that drive tumorigenesis in each genotype are likely different. Increased expression of Myc leads to apoptosis through the ARF-Mdm2-p53 tumor suppressor pathway. As a result, MYC-driven tumors often involve mutations in p19^ARF^ or p53, which render them inactive(33, 34). We have previously shown that tumors isolated from λ-*MYC* mice frequently displayed such inactivating mutations, while those from LMP2A/λ-*MYC* mice did not(22). Because LMP2A combines with Myc to increase degradation of p27^Kip1^ in our model, tumor development can occur without the necessity for inactivation of ARF-Mdm2-p53 pathway, as we have previously observed(24). We wanted to determine whether p19^ARF^ and/or p53 were frequently inactivated in tumors from p27^Super^ mice. We found that 43% (3 of 7) λ-*MYC*/p27^Super^ tumors had p19^ARF^ and/or p53 abnormalities while only 14% (1 of 7) LMP2A/λ-*MYC*/p27^Super^ tumors did (data not shown). This indicates that inactivation of the ARF-Mdm2-p53 pathway can contribute to tumor development in LMP2A/λ-*MYC*/p27^Super^ mice, but less frequently than in λ-*MYC*/p27^Super^ mice. Future studies will explore additional pathways that may differ between these two models, as well as between the LMP2A/λ-*MYC* and λ-*MYC* models in general.

Our current studies indicate the requirement that both pathways of p27^Kip1^ degradation be blocked to fully prevent the accelerated tumorigenesis driven by LMP2A. Overall, our results show that normalizing G_1_-S phase cell cycle progression by elevating levels of p27^Kip1^ both delays Myc-driven lymphoma, and offsets the contribution of LMP2A in accelerating tumor development. Our study points to both the nuclear and cytoplasmic pathways of p27^Kip1^ degradation, as well as G_1_-S phase regulation, as potential targets for developing more specific treatments of BL. Preclinical experiments testing the effectiveness of drugs targeting these pathways may uncover more effective therapies with fewer long-term side effects than the chemotherapies that are currently used.

## Materials and Methods

### Mice

The Tg6 line of Eμ-*LMP2A* transgenic mice expresses LMP2A under immunoglobulin (Ig) heavy chain promoter and intronic enhancer (Eμ), while λ-*MYC* mice overexpress human *MYC*. Both lines are in the C57BL/6 background and have been described previously(8, 19). Cks1 null (*Cks1*^−/−^) mice(28) were obtained from Steven Reed Laboratory at The Scripps Research Institute in La Jolla, CA. Mice expressing the p27^Kip1^ S10A knock-in (*Cdkn1b^tm2Jro^*)(35) were obtained from The Jackson Laboratory. Tumor mice were sacrificed when lymph node tumors could be observed externally or when mice were moribund. Animals were maintained at Northwestern University’s Center for Comparative Medicine in accordance with the university’s animal welfare guidelines.

### Tumor, spleen, and bone marrow cell isolation

Pretumor splenic B cells were purified using the Mouse Pan-B Cell Isolation Kit by StemCell Technologies. Bone marrow cells were flushed from femurs and tibia. Tumor-bearing lymph node cells were prepared as previously described(22, 24, 36).

### Flow cytometry and cell cycle analysis

To measure B cell number, purified B cells from spleens and bone marrow of 4-5 week-old mice were stained with IgM, IgD, and B220 antibodies (BD Biosciences). For cell cycle analysis, purified splenic B cells from 4-5 week-old mice were fixed in 70% ethanol and stained with propidium iodide/ribonuclease staining buffer according to manufacturer’s instructions (BD Biosciences). All flow cytometry was performed with the FACS-CantoII flow cytometer (BD Biosciences) and all results were analyzed with FlowJo software (FlowJo, LLC). Immunohistochemistry Spleens from 4-8 week-old mice and tumor bearing lymph nodes were fixed in 10% buffered formalin phosphate, stored in 70% ethanol, and embedded in paraffin. Samples were sectioned and stained with hematoxylin and eosin, anti-p27^Kip1^ (Invitrogen), or anti-B220 (BD Biosciences) antibody. Stained tissue slides were imaged using EVOS XL Core microscope.

### Immunoblots

Purified pretumor B cells or tumor cells were lysed in RIPA lysis buffer with protease and phosphatase inhibitor cocktails. Lysates were separated by sodium dodecyl sulfate polyacrylamide gel electrophoresis (Bio-Rad). Protein was transferred from the gel to a nitrocellulose membrane (Bio-Rad). Membranes were probed with anti-p27^Kip1^(Santa Cruz) or GAPDH (Abcam) primary antibodies, and then incubated with IRDye secondary antibodies (Li-Cor Biosciences). Protein bands were visualized with Odyssey Imager and analyzed with Image Studio (Li-Cor Biosciences).

### Statistical analysis

Two-tailed *t* test, survival analysis, and log-rank (Mantel-Cox) test were performed using Prism 7 (GraphPad Software). *P* < 0.05 was considered statistically significant.

## Acknowledgements

The authors thank Nanette Susmarski for timely and excellent technical assistance and members of the Longnecker laboratory for help in the completion of this study and particularly Kamonwan Fish for initiating these studies and reading the manuscript prior to submission. The authors thank the Steven Reed laboratory for the *Cks1*^−/−^ mice. The Northwestern University Mouse Histology and Phenotyping Laboratory provided histology services and are supported by NCI P30-CA060553 awarded to the Robert H. Lurie Comprehensive Cancer Center. Flow cytometry work was supported by the Northwestern University Interdepartmental Immunobiology Flow Cytometry Core Facility. This work was supported by National Institutes of Health, National Cancer Institute grant R01 CA073507 (R.L.) and Carcinogenesis Training Program grant T32CA009560 (R.P.S.). R.L. is a Dan and Bertha Spear Research Professor in Microbiology-Immunology.

## Author Contributions

R.L., R.P.S., and M.I. designed the research and wrote the manuscript. R.P.S. and M.I. performed the research and analyzed the data.

## References

1. Longnecker R, Kieff E, Cohen J. 2013. Epstein-Barr Virus, p 1898–1959. In Knipe D, Howley P (ed), Fields Virology, 6^th^ ed. Lippincott Williams & Wilkins, Philadalphia, PA.

2. Epstein MA, Achong BG, Barr YM. 1964. Virus Particles in Cultured Lymphoblasts from Burkitt’s Lymphoma. Lancet 1:702–3.

3. Babcock GJ, Decker LL, Volk M, Thorley-Lawson DA. 1998. EBV persistence in memory B cells in vivo. Immunity 9:395–404.

4. Nilsson K. 1992. Human B-lymphoid cell lines. Hum Cell 5:25–41.

5. Thorley-Lawson DA. 2015. EBV Persistence--Introducing the Virus. Curr Top Microbiol Immunol 390:151–209.

6. MacLennan IC. 1994. Germinal centers. Annu Rev Immunol 12:117–39.

7. Rickert RC. 2013. New insights into pre-BCR and BCR signalling with relevance to B cell malignancies. Nat Rev Immunol 13:578–91.

8. Caldwell RG, Brown RC, Longnecker R. 2000. Epstein-Barr virus LMP2A-induced B-cell survival in two unique classes of EmuLMP2A transgenic mice. J Virol 74:1101–13.

9. Caldwell RG, Wilson JB, Anderson SJ, Longnecker R. 1998. Epstein-Barr virus LMP2A drives B cell development and survival in the absence of normal B cell receptor signals. Immunity 9:405–11.

10. Mesri EA, Feitelson MA, Munger K. 2014. Human viral oncogenesis: a cancer hallmarks analysis. Cell Host Microbe 15:266–82.

11. Bell AI, Groves K, Kelly GL, Croom-Carter D, Hui E, Chan AT, Rickinson AB. 2006. Analysis of Epstein-Barr virus latent gene expression in endemic Burkitt’s lymphoma and nasopharyngeal carcinoma tumour cells by using quantitative real-time PCR assays. J Gen Virol 87:2885–90.

12. Chapman CJ, Zhou JX, Gregory C, Rickinson AB, Stevenson FK. 1996. VH and VL gene analysis in sporadic Burkitt’s lymphoma shows somatic hypermutation, intraclonal heterogeneity, and a role for antigen selection. Blood 88:3562–8.

13. Gregory CD, Rowe M, Rickinson AB. 1990. Different Epstein-Barr virus-B cell interactions in phenotypically distinct clones of a Burkitt’s lymphoma cell line. J Gen Virol 71 (Pt 7):1481–95.

14. Jain R, Roncella S, Hashimoto S, Carbone A, Francia di Celle P, Foa R, Ferrarini M, Chiorazzi N. 1994. A potential role for antigen selection in the clonal evolution of Burkitt’s lymphoma. J Immunol 153:45–52.

15. Muller-Hermelink HK, Greiner A. 1998. Molecular analysis of human immunoglobulin heavy chain variable genes (IgVH) in normal and malignant B cells. Am J Pathol 153:1341–6.

16. Tamaru J, Hummel M, Marafioti T, Kalvelage B, Leoncini L, Minacci C, Tosi P, Wright D, Stein H. 1995. Burkitt’s lymphomas express VH genes with a moderate number of antigen-selected somatic mutations. Am J Pathol 147:1398–407.

17. Tao Q, Robertson KD, Manns A, Hildesheim A, Ambinder RF. 1998. Epstein-Barr virus (EBV) in endemic Burkitt’s lymphoma: molecular analysis of primary tumor tissue. Blood 91:1373–81.

18. Taub R, Kirsch I, Morton C, Lenoir G, Swan D, Tronick S, Aaronson S, Leder P. 1982. Translocation of the c-myc gene into the immunoglobulin heavy chain locus in human Burkitt lymphoma and murine plasmacytoma cells. Proc Natl Acad Sci U S A 79:7837–41.

19. Kovalchuk AL, Qi CF, Torrey TA, Taddesse-Heath L, Feigenbaum L, Park SS, Gerbitz A, Klobeck G, Hoertnagel K, Polack A, Bornkamm GW, Janz S, Morse HC, 3rd. 2000. Burkitt lymphoma in the mouse. J Exp Med 192:1183–90.

20. de Alboran IM, O’Hagan RC, Gartner F, Malynn B, Davidson L, Rickert R, Rajewsky K, DePinho RA, Alt FW. 2001. Analysis of C-MYC function in normal cells via conditional gene-targeted mutation. Immunity 14:45–55.

21. Schmitt CA, McCurrach ME, de Stanchina E, Wallace-Brodeur RR, Lowe SW. 1999. INK4a/ARF mutations accelerate lymphomagenesis and promote chemoresistance by disabling p53. Genes Dev 13:2670–7.

22. Bieging KT, Amick AC, Longnecker R. 2009. Epstein-Barr virus LMP2A bypasses p53 inactivation in a MYC model of lymphomagenesis. Proc Natl Acad Sci U S A 106:17945–50.

23. Bultema R, Longnecker R, Swanson-Mungerson M. 2009. Epstein-Barr virus LMP2A accelerates MYC-induced lymphomagenesis. Oncogene 28:1471–6.

24. Fish K, Chen J, Longnecker R. 2014. Epstein-Barr virus latent membrane protein 2A enhances MYC-driven cell cycle progression in a mouse model of B lymphoma. Blood 123:530–40.

25. Kamura T, Hara T, Matsumoto M, Ishida N, Okumura F, Hatakeyama S, Yoshida M, Nakayama K, Nakayama KI. 2004. Cytoplasmic ubiquitin ligase KPC regulates proteolysis of p27(Kip1) at G1 phase. Nat Cell Biol 6:1229–35.

26. Tsvetkov LM, Yeh KH, Lee SJ, Sun H, Zhang H. 1999. p27(Kip1) ubiquitination and degradation is regulated by the SCF(Skp2) complex through phosphorylated Thr187 in p27. Curr Biol 9:661–4.

27. Fish K, Sora RP, Schaller SJ, Longnecker R, Ikeda M. 2017. EBV latent membrane protein 2A orchestrates p27(kip1) degradation via Cks1 to accelerate MYC-driven lymphoma in mice. Blood 130:2516–2526.

28. Spruck C, Strohmaier H, Watson M, Smith AP, Ryan A, Krek TW, Reed SI. 2001. A CDK-independent function of mammalian Cks1: targeting of SCF(Skp2) to the CDK inhibitor p27Kip1. Mol Cell 7:639–50.

29. Grande BM, Gerhard DS, Jiang A, Griner NB, Abramson JS, Alexander TB, Allen H, Ayers LW, Bethony JM, Bhatia K, Bowen J, Casper C, Choi JK, Culibrk L, Davidsen TM, Dyer MA, Gastier-Foster JM, Gesuwan P, Greiner TC, Gross TG, Hanf B, Harris NL, He Y, Irvin JD, Jaffe ES, Jones SJM, Kerchan P, Knoetze N, Leal FE, Lichtenberg TM, Ma Y, Martin JP, Martin MR, Mbulaiteye SM, Mullighan CG, Mungall AJ, Namirembe C, Novik K, Noy A, Ogwang MD, Omoding A, Orem J, Reynolds SJ, Rushton CK, Sandlund JT, Schmitz R, Taylor C, Wilson WH, Wright GW, Zhao EY, et al. 2019. Genome-wide discovery of somatic coding and non-coding mutations in pediatric endemic and sporadic Burkitt lymphoma. Blood doi:10.1182/blood-2018-09-871418.

30. Piccaluga PP, De Falco G, Kustagi M, Gazzola A, Agostinelli C, Tripodo C, Leucci E, Onnis A, Astolfi A, Sapienza MR, Bellan C, Lazzi S, Tumwine L, Mawanda M, Ogwang M, Calbi V, Formica S, Califano A, Pileri SA, Leoncini L. 2011. Gene expression analysis uncovers similarity and differences among Burkitt lymphoma subtypes. Blood 117:3596–608.

31. Schmitz R, Young RM, Ceribelli M, Jhavar S, Xiao W, Zhang M, Wright G, Shaffer AL, Hodson DJ, Buras E, Liu X, Powell J, Yang Y, Xu W, Zhao H, Kohlhammer H, Rosenwald A, Kluin P, Muller-Hermelink HK, Ott G, Gascoyne RD, Connors JM, Rimsza LM, Campo E, Jaffe ES, Delabie J, Smeland EB, Ogwang MD, Reynolds SJ, Fisher RI, Braziel RM, Tubbs RR, Cook JR, Weisenburger DD, Chan WC, Pittaluga S, Wilson W, Waldmann TA, Rowe M, Mbulaiteye SM, Rickinson AB, Staudt LM. 2012. Burkitt lymphoma pathogenesis and therapeutic targets from structural and functional genomics. Nature 490:116–20.

32. Abate F, Ambrosio MR, Mundo L, Laginestra MA, Fuligni F, Rossi M, Zairis S, Gazaneo S, De Falco G, Lazzi S, Bellan C, Rocca BJ, Amato T, Marasco E, Etebari M, Ogwang M, Calbi V, Ndede I, Patel K, Chumba D, Piccaluga PP, Pileri S, Leoncini L, Rabadan R. 2015. Distinct Viral and Mutational Spectrum of Endemic Burkitt Lymphoma. PLoS Pathog 11:e1005158.

33. Eischen CM, Weber JD, Roussel MF, Sherr CJ, Cleveland JL. 1999. Disruption of the ARF-Mdm2-p53 tumor suppressor pathway in Myc-induced lymphomagenesis. Genes Dev 13:2658–69.

34. Sherr CJ. 1998. Tumor surveillance via the ARF-p53 pathway. Genes Dev 12:2984–91.

35. Besson A, Gurian-West M, Chen X, Kelly-Spratt KS, Kemp CJ, Roberts JM. 2006. A pathway in quiescent cells that controls p27Kip1 stability, subcellular localization, and tumor suppression. Genes Dev 20:47–64.

36. Bieging KT, Fish K, Bondada S, Longnecker R. 2011. A shared gene expression signature in mouse models of EBV-associated and non-EBV-associated Burkitt lymphoma. Blood 118:6849–59.

